# In pursuit of biomarkers for predicting susceptibility to activity-based anorexia (ABA) in adolescent female rats

**DOI:** 10.1101/2021.10.25.465801

**Authors:** Laura K Milton, Timothy Patton, Meredith O’Keeffe, Brian J Oldfield, Claire J Foldi

**Affiliations:** Monash University, Department of Physiology, 26 Innovation Walk, Clayton, 3800 Australia; Monash Biomedicine Discovery Institute, 23 Innovation Walk, Clayton, 3800 Australia; Monash University, Department of Biochemistry and Molecular Biology, Building 77 Wellington Road, Clayton 3800 Australia; University of Melbourne, Department of Microbiology and Immunology, The Peter Doherty Institute for Infection and Immunity, Elizabeth Street Melbourne, 3000 Australia

## Abstract

Anorexia nervosa (AN) has high rates of mortality and low rates of recovery, with outcomes that worsen with illness duration. Improved early intervention strategies are required and identifying risk factors that contribute to the development of AN is critical to their implementation. The development of AN often follows a pre-existing diagnosis of anxiety disorders and obsessive-compulsive disorder and substantial genetic overlap between these conditions suggest common underlying features may predict vulnerability to AN. Moreover, patients with AN have increased levels of circulating proinflammatory cytokines, which may be involved in susceptibility to pathological weight loss considering that children with immune dysfunction have a higher risk of subsequent AN diagnoses.

Here, we used the activity-based anorexia (ABA) model to examine whether baseline levels of locomotion, anxiety-like behaviour, compulsive behaviour, and circulating immune markers predict the subsequent development of pathological weight loss in adolescent female rats. While none of these primary features were shown to differentiate rats that went on to be susceptible or resistant to weight loss in ABA, increased locomotion and anxiety-like behaviour were both associated with the *extent of* weight loss in susceptible but not resistant animals. Intriguingly, behaviour related to poor decision-making in a situation of conflict was shown to predict vulnerability to ABA. Future research using the ABA model to uncover early predictors of AN should focus on translationally relevant assays of decision-making and cognitive behaviour, dysfunction of which may not only predispose animals to ABA but may also represent an endophenotype linking anorectic, anxiety-like and compulsive behaviour.

## Introduction

Anorexia nervosa (AN) is a devastating condition with the highest mortality rate of any psychiatric disorder [1]. Up to fifty percent of patients with AN never recover [2], highlighting the urgent need for more effective therapeutic strategies. Recovery from AN becomes much less likely the longer the illness has persisted [3], therefore, early detection of symptoms and intervention are necessary for improving treatment outcomes [4]. Identifying biomarkers to help predict AN risk and/or symptom severity will enable treatment to be directed at prodromal symptoms, before weight loss becomes too severe and protracted. Both comorbid diagnoses and the expression of symptoms related to depression, anxiety and obsessive-compulsive disorder (OCD) are associated with worsened illness, poorer treatment outcomes, and increased relapse in patients with AN [5-9]. There is evidence to suggest that the development of AN often follows a pre-existing diagnosis of these comorbid conditions [10-15], presenting a potential target for early intervention strategies. However, a diagnosis of AN can also precede the development of, or be determined coincident with, comorbid psychiatric disorders [10, 13, 14], making it difficult to dissect the nature of the interaction between AN and depressive, anxious, and obsessive/compulsive symptoms.

In support of the hypothesis that anxiety and obsessional behaviour may increase susceptibility to AN in vulnerable individuals, childhood temperaments and personality traits including “picky eating”, perfectionism, harm avoidance and rigid patterns of thought and behaviour [16, 17], related to levels of anxiety and compulsiveness [10], have been associated with the subsequent development of AN. These traits often become exacerbated in the acute stage of AN and improve with weight recovery but remain elevated compared to healthy controls [10, 18], indicating that underlying personality traits are partially independent of illness status and may predispose individuals to developing AN. Moreover, there are increased diagnoses of depression, anxiety, and OCD (as well as the non-pathological symptoms of these conditions) in healthy first-degree relatives of patients with AN compared to the general population [19-21] and substantial genetic overlap between AN and generalized anxiety disorder [22], major depressive disorder [23] and OCD [24]. Taken together, these findings suggest the existence of heritable risk factors for multiple psychiatric diagnoses that cluster together and sometimes result in the development of AN. This is not surprising considering that each of these mood-related pathologies are associated with common neurobiological and behavioural phenotypes with AN [25-27].

While association of mood disorders and AN as outlined above are often established in a post hoc manner, physiological correlates should provide earlier and more definitive predictors of subsequent susceptibility to AN. In this regard, disturbances in immune function may play a causal role in the development of AN, considering that children and adolescents with autoimmune and autoinflammatory disorders are shown to have a higher risk of subsequent AN diagnoses [28] and genes associated with various immune-related phenotypes including asthma, vitiligo and type-1 diabetes are associated with AN [29]. Adults with AN have a distinct pattern of immune mediator expression, the most consistently documented of which are increased levels of the inflammatory markers RANTES (regulated upon activation, normal T cell expressed and secreted) [30], tumour necrosis factor-alpha (TNF-α) and interleukin-6 (IL-6) [31]. Pro-inflammatory cytokines produced both centrally and peripherally can have profound effects on brain function and behaviour relevant to food intake [32] and have been shown to remain elevated after weight recovery in AN patients [33]. However, the notion whereby immune markers might be used to screen for susceptibility to AN should be balanced by evidence suggesting that elevated proinflammatory cytokine expression in patients with AN is ameliorated with weight restoration [34], consistent with at least some of these changes being consequences of the conditions of chronic malnutrition typical of AN. This may include effects of a “leaky gut” induced by an anorectic state, whereby increased intestinal permeability allows antigens to traverse the intestinal wall and infiltrate systemic circulation, contributing to chronic, low-grade inflammation [35]. Intriguingly, increased levels of the same proinflammatory cytokines (IL-6 and TNF-α) are seen in patients with OCD [36] and generalised anxiety disorder is also associated with an inflammatory response that includes increased TNF-α [37]. These conditions are not associated with chronic undernutrition but, as described above, are both commonly comorbid with AN diagnoses, raising the potential that there are common inflammatory mechanisms involved in the aetiology of these disorders.

In order to determine behavioural and immune factors that contribute to the aetiology of AN, animal models are required. The most robust and widely-used animal model of AN is known as activity-based anorexia (ABA), which allows for a detailed interrogation of the biological mechanisms underlying pathological weight loss that is prohibited by the retrospective or cross-sectional nature of human studies. The ABA model pairs unhindered access to running wheels with time-limited food access and recapitulates the core features of the human condition including rapid body weight loss, voluntary hyperactivity and voluntary reductions in food intake [38-40]. We have previously shown that pathological weight loss does not occur in all rats exposed to ABA conditions, and in fact a subpopulation of adolescent female Sprague-Dawley rats remain resistant to weight loss, even when exposed to the same experimental conditions [41]. These susceptible and resistant subpopulations have been replicated in other studies investigating the neurobiology of AN in ABA rats [42] and mice [43], supporting the use of the ABA model for examining behavioural and immune phenotypes that predispose animals to pathological weight loss.

Exposure to ABA conditions is shown to increase anxiety-like behaviour on the elevated plus-maze (EPM) and the open field (OF) test [44-46], however, much like human studies, it is unclear whether this a *consequence* of weight loss or involved in vulnerability to ABA. Anxiety-like behaviour has been examined after only a single day of restricted food access in ABA mice, prior to excessive weight loss, and shown to be negatively correlated with excessive wheel running but not associated with body weight [47]. Only one study to date has examined how immune function is altered in the ABA model, with increased expression of interleukin-10 (IL-10), TNF-α and interleukin-1β (IL-1β) observed in the colonic mucosa of ABA mice compared to mice subject to the same food restriction paradigm without access to a wheel, indicating an effect specific to ABA rather than the result of limited food intake [48]. This was accompanied by elevated levels of IL-1β and its receptor (interleukin-1 receptor-1; IL-1R1) in the hypothalamus of ABA mice, although hypothalamic IL-1β expression was also increased in mice with restricted food access compared to controls. ABA exposure in this study was also shown to increase intestinal permeability, measured by circulating levels of zonulin, a plasma marker of gut barrier that modulates intercellular tight junctions [49]. Increased zonulin following ABA could therefore contribute to a “leaky gut” allowing proinflammatory molecules into circulation. It is plausible that early activation of the intestinal immune response could trigger systemic low-grade inflammation during ABA, however it remains unknown whether immune disturbances might predispose animals to developing the ABA phenotype.

In this study, we aimed to determine whether levels of anxiety-related and compulsive behaviour in rats could be used to predict susceptibility to pathological weight loss under ABA conditions, and whether differences in proinflammatory cytokine expression correlated with behaviour or the development of ABA. Our goal was to identify markers for the development of anorectic behaviour in a well-established animal model that could be used to screen at-risk individuals in order to direct early intervention strategies. This objective is in keeping with broader efforts to define predictive markers of future susceptibility to ABA in rodents and AN in humans that encompass not only behavioural but biological traits.

## Materials and methods

### Animals

Female Sprague-Dawley rats (*n*=44) were sourced from the Monash Animal Research Platform (MARP; Clayton, VIC, Australia) at 6 weeks of age with initial body weights between 120-160g. To determine whether baseline locomotor activity, anxiety-like or compulsive behaviour predicted susceptibility to weight loss in ABA, rats were tested on separate and consecutive days on the elevated plus maze (EPM), open field test (OF) and marble burying test (MBT) before exposure to ABA conditions. Blood samples were taken before behavioural testing and after recovery from the ABA paradigm to screen for proinflammatory cytokines. All experimental procedures were approved by the Monash Animal Resource Platform Ethics Committee (project ID:15171) and are detailed in **Fig 1**. A singly housed male rat was present in each experimental room to synchronise the estrous cycles of the female rats (known as the Whitten effect; [50]).

**Figure 1.**
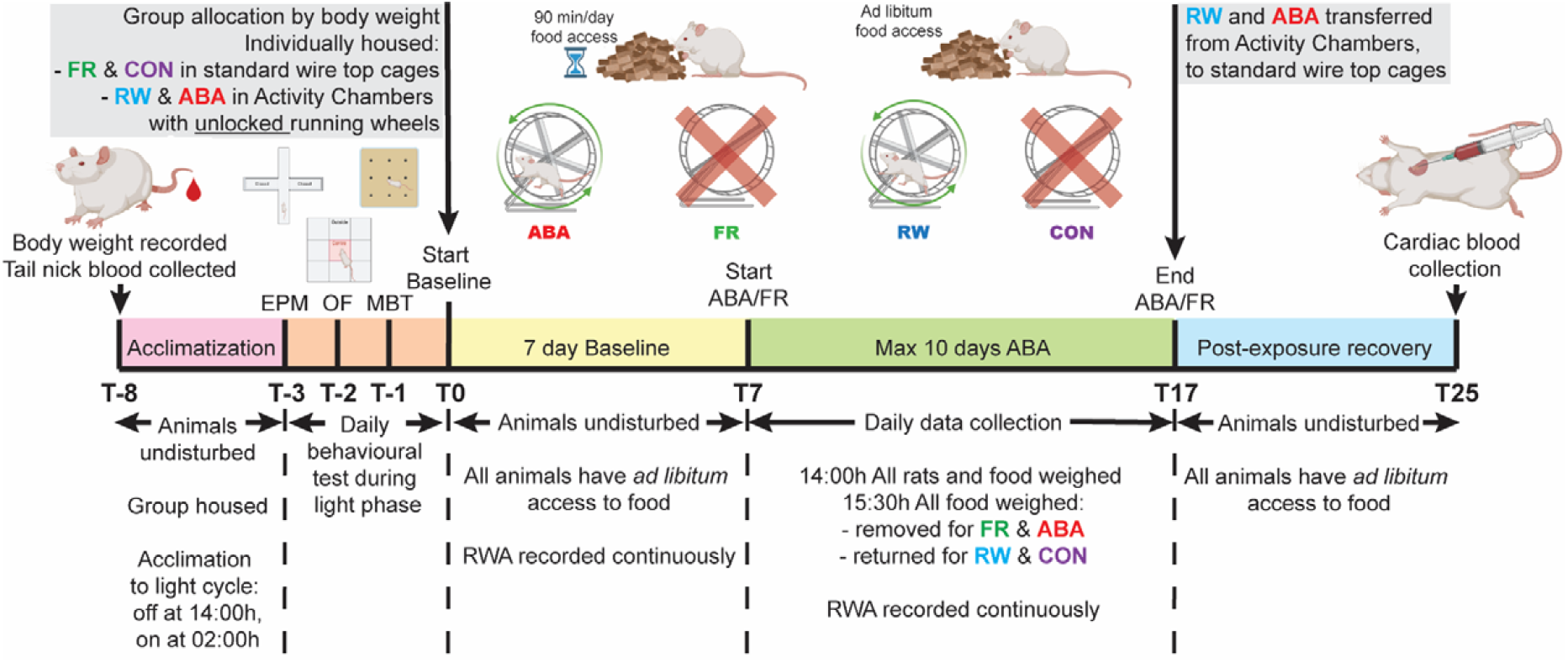
Experimental timeline. Pre-exposure blood samples were taken 8 days prior to the baseline testing period and rats were acclimated to the light cycle before maze-based behavioural tests on separate and consecutive days. The baseline testing period involved *ad libitum* food access and running activity was recorded for groups with access to wheels. Exposure to ABA conditions lasted for a maximum of 10 days, and all animals recovered with *ad libitum* access to food and no running wheels for 7 days post-exposure, followed by terminal blood collection.

### Behavioural testing

Maze-based behavioural tests were recorded with an overhead camera connected to a computer and analyzed with TopScan Lite tracking software (V 2.0; CleverSys VA, USA) and manual scoring by an observer blinded to experimental condition.

#### Anxiety-like behaviour and general locomotor activity

The elevated plus maze (EPM) consisted of an elevated 4-arm platform made of grey perspex (70 cm long × 10 cm wide × 90 cm high) with two closed (40 cm high walls) and two open arms. Rats were placed in the centre platform (10×10 cm) facing an open arm and the proportion of time spent in the closed arms relative to the open arms in each 10 min trial, was used as the primary measure of anxiety-like behaviour [51]. Frequency of head-dipping and rearing behaviour was also recorded to assess aspects of exploration. The open field (OF) test consisted of a deep open topped box (60×60×55 cm deep) in which distance travelled in each 10 min trial was used as the primary measure of locomotor activity [52] and the proportion of time spent in the aversive centre zone (middle square of a 3×3 grid; 20×20 cm) was used as a secondary measure of anxiety-related behaviour [53]. Assesment of vertical exploration was obtained during this test by recording the frequency of rearing.

#### Compulsive behaviour

The marble burying test (MBT) consisted of the same OF box described above with a 5 cm deep layer of sawdust bedding on which 9 marbles (blue glass; 8mm diameter) were placed in an even grid pattern. The “classical” measure of compulsiveness in the MBT is number of marbles buried at the end of the test duration (in this case 10 min), where a greater number indicates increased compulsive tendencies [54, 55]. However, this measure does not account for the likelihood that rats will bury and unbury marbles throughout the test, and produce a confounding ceiling effect [56], therefore the number of interactions with marbles (including burying and unburying) was used as the primary measure of compulsive behaviour in this study.

### Activity-based anorexia (ABA) and experimental controls

Four experimental groups were used to examine the influence of behavioural and immune factors on susceptibility to ABA, and rats were allocated to groups to ensure an even spread of starting body weights. Animals either underwent exposure to the ABA paradigm (ABA; *n*=20), had *ad libitum* access to food paired with access to a running wheel (RW; *n*=8), had time-limited food restriction with no wheel access (FR; *n*=8), or *ad libitum* food access and no running wheel (control; *n*=8; **see Fig 1**). Twenty rats were allocated to the ABA conditions to allow for the emergence of Susceptible and Resistant populations with adequate numbers in each. ABA and RW groups were individually housed in transparent activity wheel and living chambers (Lafayette Instruments, IN, USA; model 80859) whereas FR and control groups were singly housed in standard wire topped polypropylene cages. All groups were housed in a temperature (22-24°C) and humidity (30-50%) controlled room under a reversed 12h light/dark cycle (lights off at 1400h). Food restriction lasted for a maximum of 10 days and ABA conditions persisted until rats reached ≤80% baseline body weight, on the second day of spontaneous weight gain or for a maximum of 10 days, whichever occurred first, at which point *ad libitum* access to food was reinstated. Body weight and food intake were recorded daily between 1330h-1400h over the 10-day experimental period, with 90 min food intake also recorded at 1530h.

### Assessment of running wheel activity

Each running wheel was equipped with an Activity Wheel Counter (Lafayette Instruments, IN, USA), which was connected by USB interface to a computer running Activity Wheel Software (Lafayette instruments). Running wheel activity (RWA) was recorded in 10-minute intervals and RWA in the hour before feeding 1300-1400h was used to assess food anticipatory activity (FAA). To control for individual differences in the propensity to run, FAA was calculated as a proportion of total daily RWA and food-restriction evoked hyperactivity was calculated for individual rats as the change in running from baseline to ABA (ABA-Baseline).

### Plasma collection and cytokine analysis

Blood was collected from tail tip at 6 weeks of age, 5 days prior to the commencement of behavioural testing. Approximately 350 ul of whole blood was collected into EDTA coated tubes (Microvette; Brand), which were centrifuged for 10 min (8 000 rpm, 4 °C) within 30 min of collection and plasma separated and stored at -80 °C until use. Following exposure to ABA conditions, 7 days of *ad libitum* food access and body weight recovery to >100% baseline, blood was collected via cardiac puncture. A custom rat multi-analyte LEGENDplex bead-based immunoassay kit was used to examine cytokine concentrations in plasma samples (LEGENDplex; BioLegend, CA, USA) that targeted six cytokines concurrently (IL-6, IL-10, IL-4, IL-1β, TNF-α, RANTES). These analytes were selected based on their previously reported elevation in human AN patients and/or ABA mice. Plasma samples were screened with the LEGENDplex assay kit as per manufacturer’s instructions, and the readout measurement acquired using a Fortessa X-20 flow cytometer (Becton Dickinson (BD); Franklin Lakes, NJ, USA). Data were analysed using LEGENDplex Data Analysis Software (v8.0; BioLegend).

### Statistical analyses

Except otherwise noted all statistical analysis was performed in GraphPad Prism 8.0 (GraphPad Software, San Diego, CA). Significance for all tests was set at *p*<0.05. A variety of statistical analyses were used determined by the type of data and number of groups: One- and Two-way ANOVA, followed by post-hoc multiple comparisons (Tukey’s or with a Bonferroni correction) when applicable; independent samples t-test; Pearson’s correlation; and linear regression. Details of each individual analysis and complete statistical results can be found in Supplementary Results.

### Exclusions

One rat exposed to ABA showed an abnormal body weight loss trajectory and one rat exposed to FR was unable to maintain body weight >80% of baseline; data collected from both these animals was excluded from all analyses. Concentrations of IL-10, TNF-α and IL-1β were not detected at levels high enough to be reliable using the LEGENDplex assay so these analytes were excluded from all analyses. Any pre- and post-exposure pair of RANTES, IL-4 or IL-6 samples that was incomplete was excluded from analyses.

## Results

### Effects of individual components of the ABA paradigm on weight maintenance and feeding

Of rats exposed to ABA conditions (**Fig 2A**), 9/19 (47%) were resistant to body weight loss (**Fig 2B**), and while weight loss was negatively correlated with food intake for Resistant animals, in line with all three control groups (all *p*s<.0001), there was no such association for rats Susceptible to ABA (*p*=.1610; **Fig 2C**). Moreover, the body weight loss trajectories for Resistant rats plateaued similarly to rats without access to a running wheel (FR; **Fig 2D**), whereas Susceptible rats lost significantly more body weight on average per day than both Resistant and FR rats (both *p*s<.0001; **Fig 2E**). Access to a running wheel alone did not produce significant weight loss (RW, *p*=.1446; **Fig 2F**) but did increase food intake compared to control rats (*p*=.0003; **Fig 2G** and **I**) presumably to compensate for increased energy expenditure from wheel running. Much like the pattern of body weight loss, Resistant rats ate a similar amount of food to FR rats (*p*=.9982; **Fig 2G** and **H**) whereas Susceptible rats ate significantly less than both other food restricted groups (both *p*s<.0001; **Fig 2G** and **H**).

**Figure 2.**
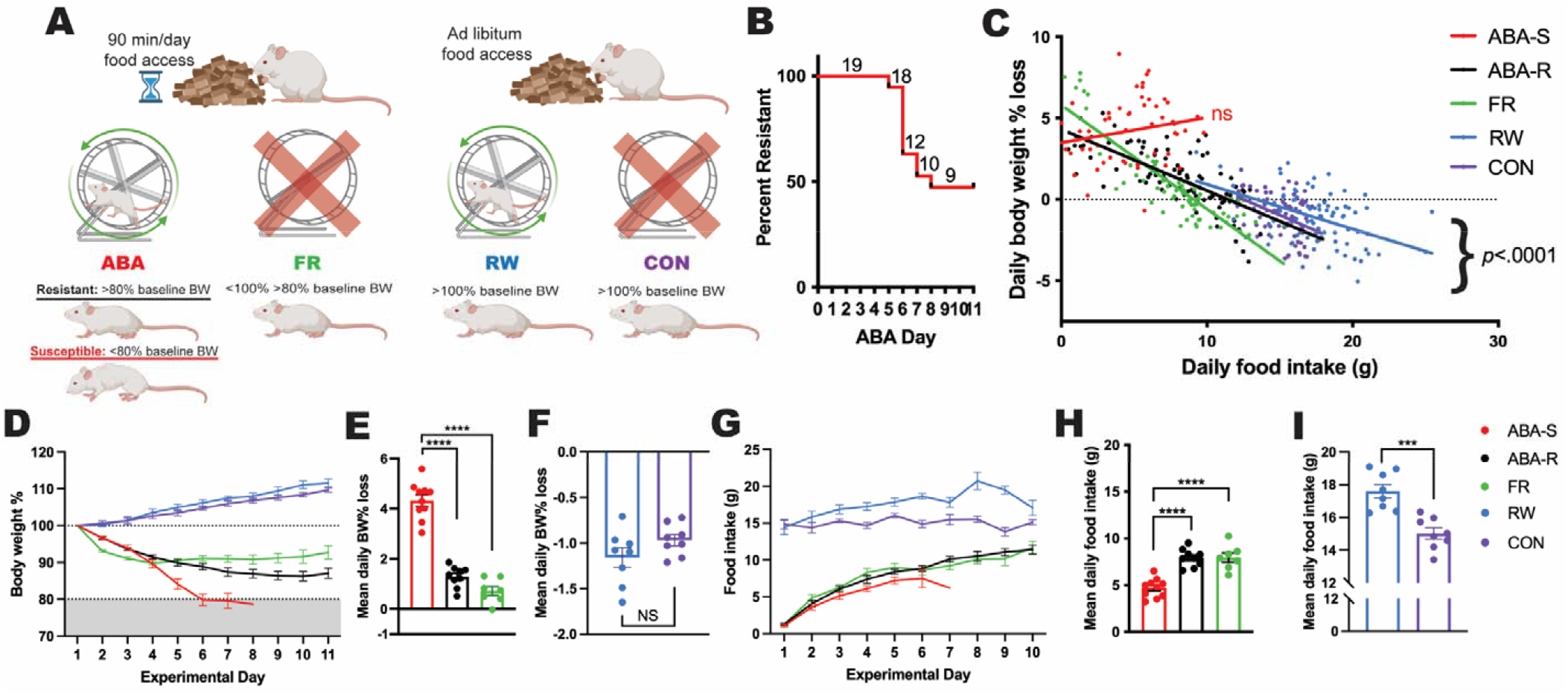
Body weight and food intake following exposure to individual or combined components of the ABA paradigm. **A)** Schematic showing the four experimental treatments and five resulting groups: ABA-S, rats that were Susceptible to ABA; ABA-R, rats that were Resistant to ABA; FR, sedentary rats with food restriction; RW, *ad libitum* fed rats with access to running wheels; CON, *ad libitum* fed sedentary controls. **B)** Plot of animals remaining in the ABA paradigm over time: 9/19 (47%) rats were resistant to ABA. **C)** Linear regression with line of best fit and each individual daily data point for all animals (full results in Supplementary Table 1). Daily food intake and daily body weight % loss were significantly negatively correlated for ABA-R (*r*=.7248), FR (*r*=.8712), RW (*r*=.4486) and CON (*r*=.4434; all *p*s<.0001), whilst there was no relationship between these variables for ABA-S (*p*=.1610). Daily body weight % **(D)** and daily food intake **(G)** over the experimental period; group mean SEM. Mean daily body weight % loss **(E** and **F)** and food intake **(H** and **I)**; individual animals (circles) and group mean SEM. **E** and **H)** One-way ANOVA (results in Supplementary Table 1) followed by Tukey’s post-hoc multiple comparisons. ABA-S rats lost significantly more body weight % **(E)** and ate significantly less **(H)**, on average, per day than both ABA-R and FR groups (all *p*s<.0001). **F** and **I)** Independent samples t-test (results in Supplementary Table 1). **F)** Mean daily body weight % loss was not significantly different for RW and CON (*p*=.1446). **I)** RW ate significantly more, on average, per day than CON (*p*=.0003). NS = not significant *p*>.05, ^***^*p*<.001, ^****^*p*<.0001.

### Running wheel activity predicts susceptibility to weight loss under ABA conditions

Consistent with our previous findings [41], running activity in the two days prior to food restriction predicted susceptibility to pathological weight loss in ABA, and food restriction-evoked hyperactivity did not occur in animals resistant to weight loss (all *p*s<.0001; **Fig 3B**). Food restriction-evoked hyperactivity was especially pronounced in Susceptible rats during dark phase running (all *ps*<.0063), but also occurred in the light phase (*p*=.0156; **Fig 3C**). This shift toward light phase running in Susceptible animals was evident by the third day of ABA, with the light-phase activity peak constituting almost all running activity in Susceptible rats by ABA Day 7 (**Fig 3D**). Resistant rats also increase light-phase running during ABA, but preferentially within the hour preceding food access (*p*<.0001; **Fig 3E**), a deliberate and motivated increase in anticipation of food (FAA). On average, there were no differences between groups during baseline running and Resistant rats ran at levels similar to RW rats, whereas Susceptible rats show marked hyperactivity during ABA compared to both other groups (both *p*s<.0001; **Fig 3F**). When we examined the change in running activity elicited by ABA in individual rats to account for baseline variability, hyperactivity remained specific for Susceptible rats compared to Resistant (*p*=.0022) and RW rats (*p*=.0008; **Fig 3G**). Interestingly, daily running was positively correlated with body weight for RW (*p*=.0012) but not Resistant rats (*p*=.1476; **Fig H**) and, conversely, associated with greater food intake for Resistant rats (*p*<.0001) but not RW rats (*p*=.0774; **Fig 3I**) suggesting that while overall running did not differ between these groups, the relationship between running and other measures of energy balance did. The aspect of running that was specifically increased in Resistant rats, FAA, was elevated over Susceptible and RW levels as both a percentage of running activity during ABA (**Fig 3J**) and change from baseline levels (all *p*s<.0019; **Fig 3K**), and was inversely associated with body weight loss in Resistant animals (*p*<.0001; **Fig 3L**). FAA is considered to be a motivated behaviour that occurs in periods of food scarcity to increase the likelihood of finding food. It is therefore noteworthy that while FAA was positively associated with food intake for Resistant (*p*<.0001) and RW (*p*=.0027) animals, this association was absent in rats susceptible to ABA (*p*=.9293; **Fig 3M**).

**Figure 3.**
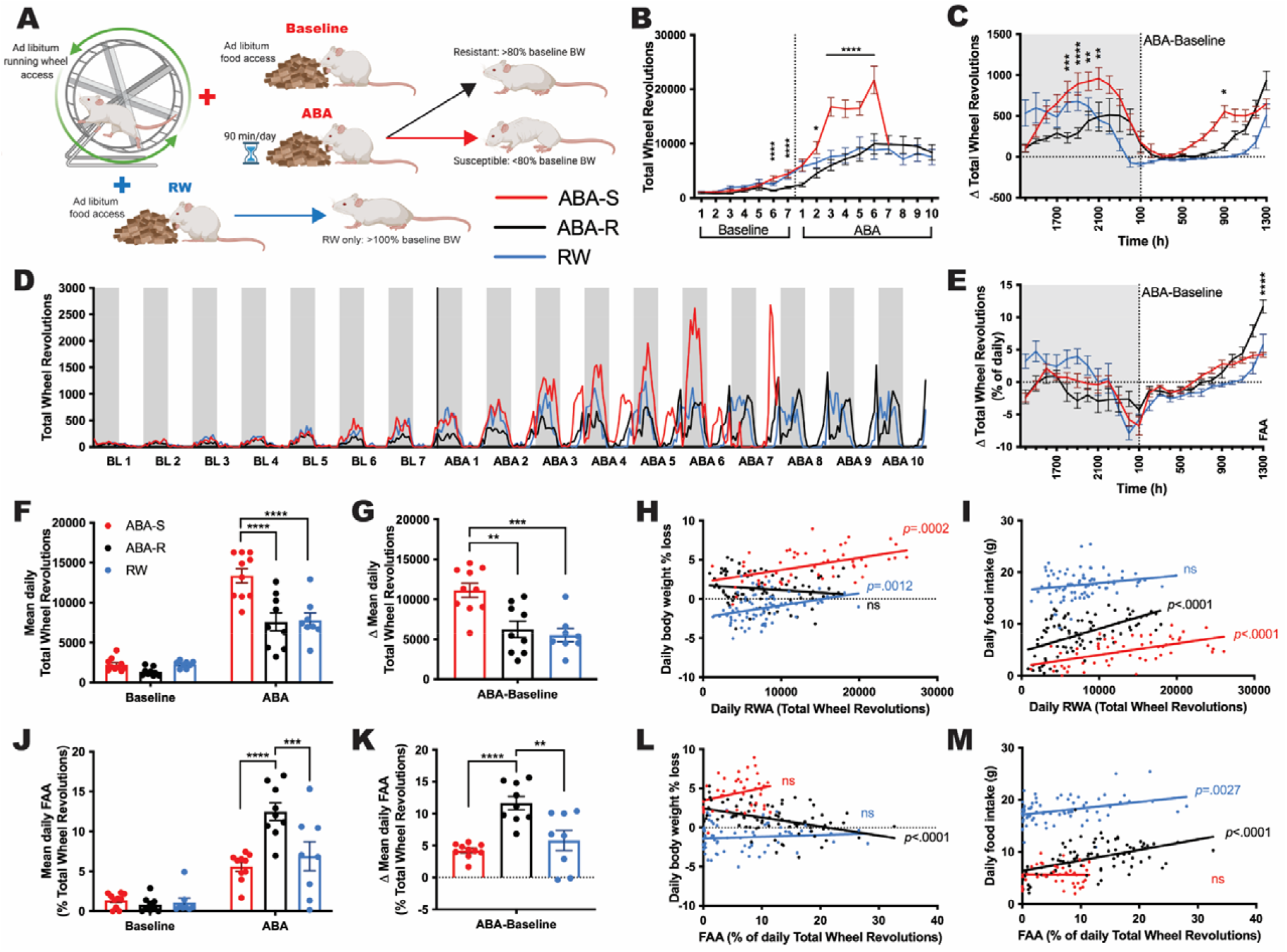
RWA for rats with wheel access either maintained on *ad libitum* feeding (RW) or combined with food restriction (ABA). **A)** Schematic showing the three experimental groups: Susceptible to ABA (ABA-S), Resistant to ABA (ABA-R), and those with access to running wheels but maintained on *ad libitum* feeding (RW). Group mean SEM of daily RWA across the entirety of the experiment **(B)**, change in mean hourly RWA across the dark (grey) and light (white) phases from baseline to ABA periods as absolute wheel revolutions **(C)**, and as a proportion of total daily RWA **(E). D)** Hourly RWA across dark (grey) and light (white) phases for the entirety of the experiment; group mean without error shown for clarity. **B, C, E)** Two-way RM ANOVA (results in Supplementary Table 2) followed by post-hoc multiple comparisons with a Bonferroni correction were performed <u>only on the ABA groups</u>, RW is graphed solely for visual comparison. **B)** Overall, ABA-S rats had significantly greater daily RWA than ABA-R rats across both the Baseline (*p*=.0101) and ABA (*p*<.0001) periods. <u>Baseline</u>: Future ABA-S rats ran significantly more on baseline days 6 and 7 than future ABA-R rats (both *p*s<.0001). <u>ABA</u>: ABA-S rats had significantly greater RWA than ABA-R rats on ABA day 2 (*p*=.014) and days 3-6 (all *p*s<.0001). **C)** ABA-S rats had a significantly greater increase in hourly RWA from baseline to ABA than ABA-R rats from 1800h-2100h (all *p*s<.0063), and 0900h (*p*=.0156). **E)** ABA-R rats had a significantly greater increase in proportional mean hourly RWA from baseline to ABA than ABA-S rats in the penultimate hour before food access (1300h; *p*<.0001). Mean daily RWA **(F)** and proportional FAA **(J)** during baseline and ABA phases, and change in mean daily RWA **(G)** and proportional FAA **(K)** from baseline to ABA; individual animals (circles) and group mean SEM. **F** and **J)** Two-way RM ANOVA (results in Supplementary Table 2) followed by post-hoc multiple comparisons, Tukey’s for between groups or with a Bonferroni correction for between phase. **G** and **K)** One-way ANOVA (results in Supplementary Table 2) followed by Tukey’s post-hoc multiple comparisons. **H, I, L, M)** Linear regression with line of best fit and each individual daily data point for all animals (full results in Supplementary Table 2). **F)** ABA-S rats ran significantly more, on average, per day in the ABA phase than both ABA-R and RW rats (both *p*s<.0001). **G)** The magnitude of increase in mean daily RWA from baseline to ABA was significantly greater in ABA-S rats than both ABA-R (*p*=.0022) and RW (*p*=.0008) rats. **H)** Daily RWA and Daily body weight % loss were significantly positively correlated for ABA-S (*r*=.4970, *p*=.0002) and RW (*r*=.3553, *p*=.0012) animals, with no relationship between these variables for ABA-R animals (*p*=.1476). **I)** Daily RWA and Daily food intake were significantly positively correlated for ABA-S (*r*=.5574, *p*<.0001) and ABA-R (*r*=.5508, *p*<.0001) animals, whilst there was no relationship between these variables for RW animals (*p*=.0774). **J)** ABA-R rats had significantly greater FAA in the ABA phase than both ABA-S (*p*<.0001) and RW (*p*=.0002) rats. **K)** The magnitude of increase in mean daily FAA from baseline to ABA was significantly greater in ABA-R rats than both ABA-S (*p*<.0001) and RW (*p*=.0019) rats. **L)** Daily FAA and Daily body weight % loss were significantly negatively correlated for ABA-R (*r*=.4569, *p*<.0001) while there was no relationship between these variables for either ABA-S (*p*=.0666) or RW (*p*=.4237) animals. **M)** Daily FAA and Daily food intake were significantly positively correlated for both ABA-R (*r*=.4876, *p*<.0001) and RW (*r*=.3484, *p*=.0027) animals, but there was no relationship between these variables for ABA-S animals (*p*=.9293). ^*^*p*<.05, ^**^*p*<.01, ^***^*p*<.001, ^****^*p*<.0001.

### Measures of anxiety-like and compulsive behaviour do not predict subsequent susceptibility to ABA

The duration of time spent in different zones of the EPM (**Fig 4A**) did not differentiate between rats that went on to be susceptible or resistant to ABA (*p*=.9979; **Fig 4B**); however, rats that went on to be susceptible to ABA entered and exited zones more often than rats that went on to be resistant to ABA (*p*=.0053; **Fig 4C**), exemplified by more frequent crosses into the centre zone (*p*=.0355), without an overall increase locomotor activity (*p*=.1215; **Fig 4D**). The number of exploratory rears (*p*=.6048; **Fig 4E**) or head dips in the EPM (*p*=.2709; **Fig 4F**) did not differ between rats that went on to be susceptible or resistant to ABA, however, the primary measure of anxiety-like behaviour on the EPM, duration spent in the open arms, was correlated with the extent of weight loss in ABA *for Susceptible rats only* (*p=*.0421; **Fig 4G**). Similarly, while measures of locomotion, anxiety-like or exploratory behaviour in the OF (**Fig 4H**) did not predict susceptibility to ABA (distance travelled *p*=.8629, duration in centre *p*=.5724, entries into centre *p*=.6799, rearing *p*=.7875; **Fig 4I-L)**, locomotor activity was correlated with the extent of weight loss in Susceptible (*p=*.0091) but not Resistant (*p*=.4891) rats (**Fig 4M**). Of note, given that RWA at baseline was predictive of ABA susceptibility but general locomotor activity was not, there was also no relationship between total distance travelled in the OF and baseline RWA (Susceptible, *p*=.1538; Resistant, *p*=.5278; **Fig 4N**). Moreover, while there was a trend toward increased interactions with marbles in the MBT (**Fig 4O**) for rats that went on to be resistant to ABA (*p*=.0551; **Fig 4P**), this measure was not correlated with running wheel activity at baseline or during ABA (all *p*s>.1873; **Fig 4Q**) but was positively associated with food intake for Resistant (*p=*.0305) but not Susceptible (*p*=.1319) rats (**Fig 4R**). Considering that Resistant rats ate significantly more than Susceptible rats, this suggests that interactions with marbles may be less reflective of compulsive behaviour and perhaps represents increased exploration directed toward novelty (i.e. the introduction of marbles or food hoppers to the environment).

**Figure 4.**
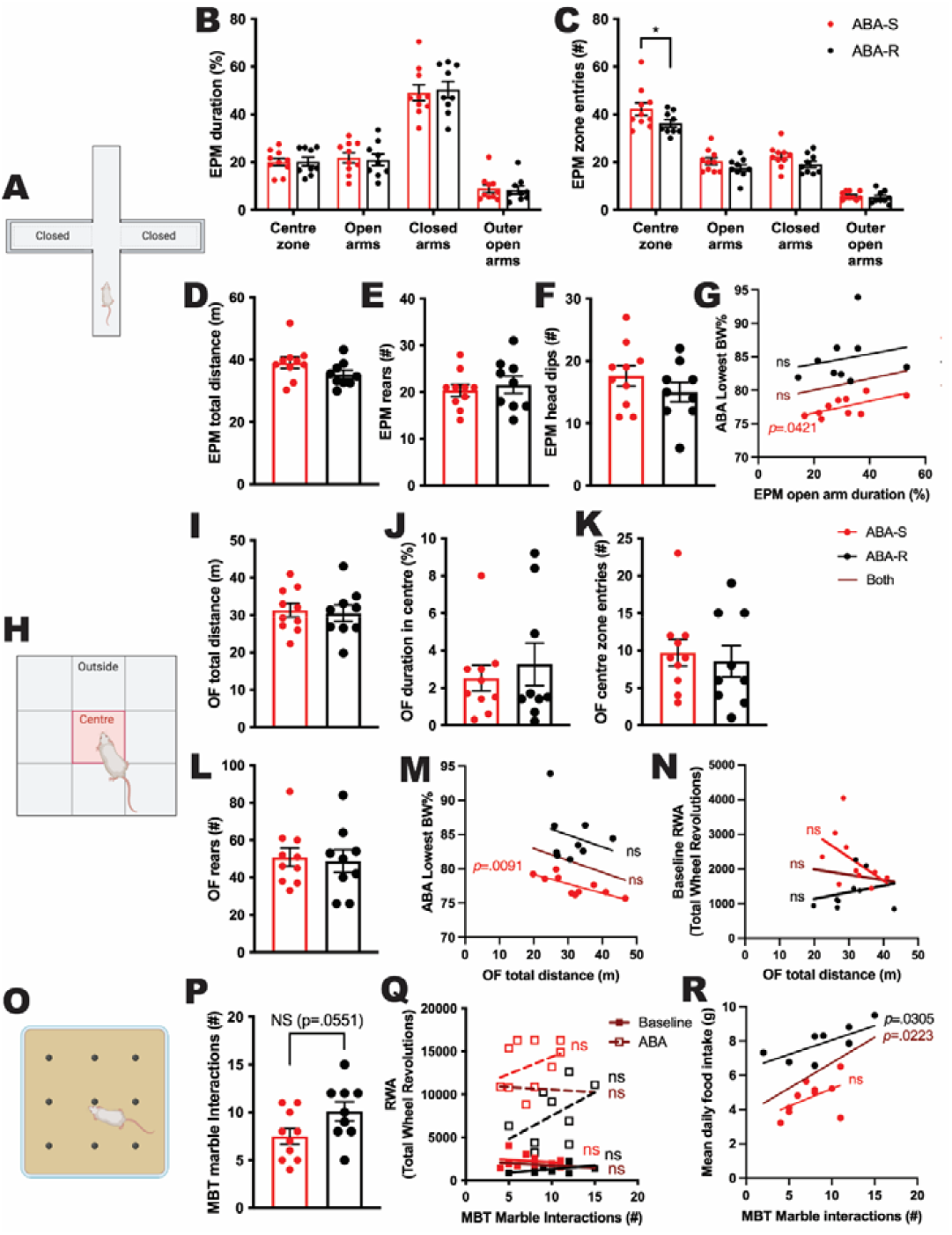
Pre-exposure behavioural test results and correlations with ABA outcomes. **A)** Schematic of the EPM; outer open arms are not shown but constituted the end 1/3 of each open arm. Bar graphs show group mean SEM with individual animals (circles) for animals that went on to be Susceptible to ABA (ABA-S) or Resistant to ABA (ABA-R). **B** and **C)** Two-way RM ANOVA (results in Supplementary Table 3). **B)** Duration (%) spent in each of the EPM zones did not differ significantly between groups. **C)** Number of entries into each of the EPM zones. Future ABA-S rats made significantly more entries into the centre zone than future ABA-R rats (*p*=.0355). **D-F, I-L, P)** Independent samples t-test (results in Supplementary Table 3). Neither the total distance (**D**), number of exploratory rears (**E**), nor number of head dips (**F**) on the EPM differed between groups (all *p*s>.1215). **G, M, N, Q, R)** Linear regression with line of best fit and all individual animal data points; line of best fit for the combined ABA group is also shown (full results in Supplementary Table 3). **G)** The duration of time spent in the open arms of the EPM was significantly positively correlated for ABA-S rats (*r*=.6495, *p*=.0421), whilst there was no relationship between these variables for ABA-R (*p*=.5801) nor the combined ABA group (*p*=.4248). **H)** Schematic of the OF. None of the OF outcome measures differentiated between rats that went on to be Susceptible or Resistant to ABA (all *p*s>.5724): **I)** Total distance; **J)** Centre duration (%); **K)** Centre zone entries; **L)** Exploratory rears. **M)** OF total distance was significantly negatively correlated with subsequent lowest body weight % during ABA for ABA-S rats (*r*=.7706, *p*=.0091), but not for ABA-R rats (*p*=.4891) or the combined ABA group (*p*=.2753). **N)** There was no relationship between OF total distance and mean daily Baseline RWA for any group (all *p*s>.1538). **O)** Schematic of the MBT. **P)** The number of interactions with marbles was not significantly different between groups (*p*=.0551), was not significantly correlated with mean daily RWA during either Baseline or ABA for any group (**Q;** all *p*s>.1873), but was significantly positively correlated with 90-minute food intake during ABA **(R)** for ABA-R (*r*=.7147, *p*=.0305) and the combined ABA group (*r*=.5205, *p*=.0223), but not for ABA-S (*p*=.1319).

### Expression of pro-inflammatory cytokines does not predict susceptibility to ABA

Expression of IL-10, IL-1β and TNF-α was below the assays limit of detection in a large proportion of samples tested, precluding reliable comparisons between groups (data not shown). Expression of RANTES (**Fig 5**_**B1-B3**_), IL-4 (**Fig 5**_**C1-C3**_) and IL-6 (**Fig 5**_**D1-D3**_) was not significantly different between groups prior to exposure to the various experimental conditions, was not systematically altered by access to a running wheel or restricted food access and did not predict susceptibility or resistance to pathological weight loss under ABA conditions. However, baseline expression of IL-6 was positively associated with change in body weight for Resistant but not Susceptible rats (*p=*.0305 and *p*=.4627 respectively; **Fig 5E**), and the association between IL-6 and running activity was in opposite directions for Susceptible and Resistant rats, with increased IL-6 associated with higher running activity for Susceptible (*p*=.0404) but lower running activity for Resistant rats (*p=*.0386; **Fig 5F**). Moreover, the extent to which ABA elicited a change in IL-6 expression was associated with food intake only for Susceptible (*p=*.0128) but not Resistant (*p*=.9219) rats (**Fig 5G**). Finally, the change in IL-4 expression elicited by ABA was associated with increased weight loss for Susceptible (*p=*.0173) but not Resistant (*p*=.8211) rats (**Fig 5H**).

**Figure 5.**
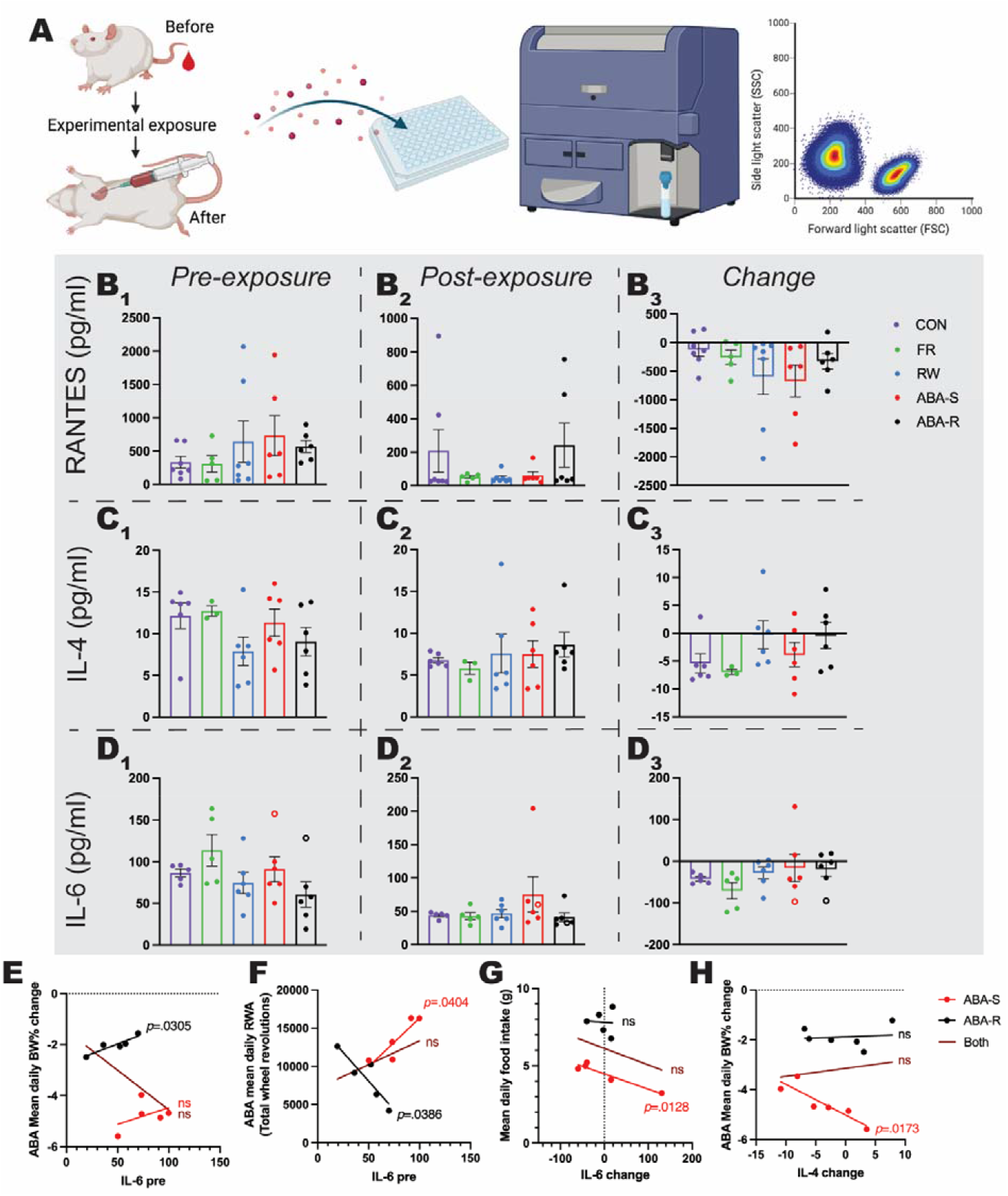
Concentration of proinflammatory cytokines before and after exposure to ABA conditions. **A)** Schematic of blood collection and flow cytometry timeline. **B-D)** Plasma cytokine concentration (pg/mL) collected from rats before experimental exposure **(1)**, at the cessation of the experiment **(2)** and the change from before to after **(3)** for RANTES **(B)**, IL-4 **(C)**, and IL-6 **(D)**. Welch’s One-way ANOVAs revealed no significant differences between groups for any cytokine (all *p*s>.0581; results in Supplementary Table 4). **E-G)** Linear regression with line of best fit and all individual animal data points (full results in Supplementary Table 4); <u>NOTE</u>: two outlying IL-6 pre-exposure values (1 from each ABA subgroup; open circles in **D**), and the corresponding change values, were excluded from linear regression analyses leaving n=5 in each group. **E)** Pre-exposure levels of IL-6 were significantly positively correlated with mean daily body weight % change in ABA, such that higher pre-exposure IL-6 was associated with less weight loss, for ABA-R (*r*=.9128, *p*=.0305), but not ABA-S (*p*=.4627) nor the combined ABA group (*p*=.1396). **F)** Pre-exposure levels of IL-6 were significantly correlated with mean daily RWA during ABA for both ABA subgroups, although in opposite directions; positive for ABA-S (*r*=.8947, *p*=.0404) and negative for ABA-R (*r*=.8978, *p*=.0386). Consequently, there was no significant relationship between these variables for the combined ABA group (*p*=.2617). **G)** The change in IL-6 from pre- to post-exposure was significantly negatively correlated with mean 90-minute food intake during ABA for ABA-S (*r*=.9513, *p*=.0128), but not ABA-R (*p*=.9219) nor the combined ABA group (*p*=.3963). **H)** The change in IL-4 from pre- to post-exposure was significantly negatively correlated with mean daily body weight % change in ABA for ABA-S (*r*=.8907, *p*=.0173), but not ABA-R (*p*=.8211), nor the combined ABA group (*p*=.7101).

## Discussion

Mood-related comorbidities and immune dysregulation both contribute to increased risk and severity of AN in human patients. Here, we show that neither the behavioural correlates of anxiety, locomotion and compulsivity nor circulating inflammatory markers that are associated with AN contribute to susceptibility to pathological weight loss in the ABA rat model. Contrary to expectation, classical measures of anxiety-like behaviour or general locomotor activity did not differentiate between rats that went on to be susceptible or resistant to ABA. However, specific aspects of the ABA microstructure were differentially associated with behavioural phenotypes in Susceptible versus Resistant rats, providing new insight into why pathological weight loss occurs in some animals exposed to ABA conditions but not others. For example, increased anxiety-like behaviour on the EPM and hyperlocomotion in the OF were associated with a greater *extent* of weight loss in ABA for Susceptible but not Resistant rats, suggesting that these behavioural features contribute to increased weight loss in Susceptible individuals. This is consistent with evidence that childhood or adolescent anxiety disorder symptoms are associated with lower BMI in individuals with AN [57].

Moreover, rats that went on to be susceptible to weight loss in ABA demonstrated a significant increase in the number of crossings between zones on the EPM, which was driven by more frequent centre crossings. The centre of the EPM is considered to be a point of choice or decision-making, from which animals engage in high levels of risk assessment [58]. Thus, if an animal is particularly indecisive about which arm to explore, the result is increased centre platform activity [59], which has been associated with inefficient responding in an operant conditioning task [60]. This suggests that susceptibility to weight loss in ABA in the present study was influenced by cognitive function, specifically poor decision-making. Deficits in cognitive capacities are common to patients both currently ill with and weight-recovered from AN [61-64], and cognitive function is impaired in rats after exposure to ABA [65]. We have recently shown a neurobiological link between pathological weight loss in ABA and cognitive flexibility using touchscreen-based assays of operant responding [66]. There is also evidence that cognitive dysfunction is a risk factor for general psychopathology in adolescents [67], and may therefore represent an endophenotype linking the comorbid development of mood disorders and AN. Future research should use translational assays of cognitive behaviour, such as those based in touchscreen technology, prior to exposure to ABA conditions in order to further interrogate this association.

This study also recapitulated our previous finding [41] that susceptibility to ABA could be predicted from baseline running wheel activity in the two days prior to initiation of food restriction, and built upon this to show that general locomotor activity was correlated with the extent of weight loss in Susceptible but not Resistant animals. However, the proportion of rats that were Resistant to developing the ABA phenotype in this study was 47% - a marked increase from the ∼30% previously reported [41, 68]. It is likely that the additional handling required for maze-based behavioural testing is responsible for this increase, considering that repeated postnatal handling reduces vulnerability to ABA in adult rats [69]. It was also evident from the present study that locomotor and running activity were not correlated for any group of animals that had access to wheels. This discrepancy highlights the fact that general locomotion and activity in the running wheel are distinct behaviours, likely underpinned by different neurochemical and motivational processes. While general locomotor activity is a relatively constant behaviour, wheel running is a deliberate and driven behaviour requiring a conscious decision to engage in, and effort to perform [70]. Wheel running in ABA is also thought to be compulsive [71], however in the present study, RWA was not associated with any measure on the MBT and in fact there was a trend toward increased “compulsive” marble burying for rats that went on to be *resistant* to ABA. This indicates that either compulsive behaviour develops coincident with access to the wheel during ABA, rather than predisposes animals to pathological weight loss, or that the MBT is not a reliable measure of compulsivity in rats, but perhaps relates to investigative drive [72]. In support of the latter, marble burying was correlated with increased food intake for Resistant but not Susceptible rats, which may reflect an increased willingness to explore novel/changing environments (in this instance increasing approach to the food hopper and subsequent food intake when available).

With respect to how circulating proinflammatory cytokines might predict susceptibility to ABA, there was no consistent change in expression of RANTES, IL-4 or IL-6 that was associated with any aspect of the ABA paradigm when sampled at the baseline, pre-exposure state. This is perhaps unsurprising considering changes in cytokine expression normally follow a “challenge”, however, it was interesting to see that IL-6 expression prior to exposure to ABA was associated with *increased* running activity for Susceptible rats, but *decreased* running activity for Resistant rats, highlighting a divergence between these populations. It is clear that the intensity and duration of exercise are key factors in determining changes in inflammatory cytokine expression, and that exercise-mediated inflammation presents a different effect depending on the tissue type examined. For example, chronic or exhaustive exercise in rats both increases cytokine concentrations in skeletal muscle and reduces concentrations in adipose tissue [73, 74], whereas moderate exercise reduces expression of inflammatory cytokines in skeletal muscle [75] and in serum [76]. Surprisingly, running activity appeared not to alter markers of inflammation in the present study, with no significant changes in circulating cytokines elicited by either normal or excessive running behaviour. It is plausible that any changes in expression related to running activity may have been masked by opposing changes in tissue-specific release. Moreover, the effects of caloric restriction on pro-inflammatory cytokine production is similarly tissue-specific [77] but both intermittent and chronic calorie restriction paradigms are shown to reduce serum IL-6 among other cytokines compared to *ad libitum* feeding [78]. There were no significant changes in cytokine expression elicited by food restriction in the present study, but the change in IL-6 expression following exposure to ABA was associated with reduced food intake only for Susceptible rats. It should be noted that post-exposure blood samples were collected after body weight recovery to at least baseline levels, which also included a period of *ad libitum* food access for FR and ABA rats and no running wheels for RW and ABA rats. Considering the points above, and that elevated inflammatory markers seen in patients with AN have recently been shown to normalise with body weight recovery [34], it may be that a dysregulated inflammatory state is confined to the acute stage of anorexia in both humans and rats.

Despite some interesting hints that aspects of decision-making and anxiety-related behaviour could be used to predict susceptibility to pathological weight loss in the ABA model, this study did not identify a set of reliable behavioural or inflammatory features that predispose animals to ABA. It may be premature, however, to exclude such predisposing behaviours on the basis of the tests performed. These may lack the sensitivity or acuity to unveil underlying differences that will align with susceptibility to ABA. Alternatively, there may be no single behavioural or immune-related measure that predicts susceptibility to ABA, but rather a raft of small differences that independently fail to predict susceptibility but, in concert, comprise a cumulative burden which promotes susceptibility to ABA. This would fit with the difficulties in finding predictive markers for AN in humans and also explain the deficiencies of current treatments that often target only a subset of AN pathologies. It should be recognized that despite our intention to incorporate the most appropriate behavioural tests that may highlight a predisposition, this list is not exhaustive and that other predictive tests may be developed and utilised in the future, particularly with respect to cognitive mediators of susceptibility to ABA. It may also be the case that traditional behavioural testing, requiring extensive handling and experimenter intervention, is not the most appropriate means by which to examine predisposing behaviours for ABA, considering that repeated handling is known to alter susceptibility to ABA [69]. It is now possible with novel technologies to examine naturalistic behaviour in rodent models *without* experimenter intervention, for example using fully automated testing systems that rely on RFID technology to gate access of individual animals to testing chambers [79, 80]. Combining these automated systems with translationally relevant assays of cognition using touchscreens as described above offers a powerful tool for future investigations of the cognitive and behavioural predictors of ABA.

## Acknowledgements

The authors acknowledge the use of the facilities at Monash FlowCore and financial support from a Research Training Program (RTP) stipend (LKM) and a Rebecca L Cooper Medical Research Foundation Project Grant (CJF). Figures were created with resources from Biorender.com.

---

**Supplementary Table 1.** Statistical test details and results for data presented in Figure 2.

| **Figure** | **Statistical test** | **Main analysis result** | **Post-hoc multiple comparisons** |
| --- | --- | --- | --- |
| **2B** | Linear regression | ABA-S: $r=.1935$, $R^2=.03816$, $p=.1610$<br>ABA-R: $r=.7248$, $R^2=.5254$, **$p<.0001$**<br>FR: $r=.8712$, $R^2=.7509$, **$p<.0001$**<br>RW: $r=.4486$, $R^2=.2012$, **$p<.0001$**<br>CON: $r=.4434$, $R^2=.1966$, **$p<.0001$** | |
| **2E** | One-way ANOVA | $F(2, 23)=101.3$, **$p<.0001$** | ABA-S > ABA-R, **$p<.0001$**<br>ABA-S > FR, **$p<.0001$** |
| **2F** | Independent samples t-test | $t(14)=1.545$, $p=.1446$ | |
| **2H** | One-way ANOVA | $F(2, 23)=27.29$, **$p<.0001$** | ABA-S < ABA-R, **$p<.0001$**<br>ABA-S < FR, **$p<.0001$** |
| **2I** | Independent samples t-test | $t(14)=4.846$, **$p=.0003$** | |

**Supplementary Table 2.** Statistical test details and results for data presented in Figure 3.

| **Figure** | **Statistical test** | **Main analysis result** | **Post-hoc multiple comparisons** |
| --- | --- | --- | --- |
| **3B** | Baseline: Two-way RM ANOVA<br><u>ONLY INCLUDES ABA-S and ABA-R</u> | Time $F(6, 102)=20.24$, $p<.0001$<br>Outcome $F(1, 17)=8.378$, $p=.0101$<br>Interaction $F(6, 102)=6.850$, $p<.0001$ | Day 6: ABA-S > ABA-R $p<.0001$<br>Day 7: ABA-S > ABA-R $p<.0001$ |
| **3B** | ABA: Mixed-model ANOVA (due to missing values from ABA-S rats removed from the experiment over time)<br><u>ONLY INCLUDES ABA-S and ABA-R</u> | Time $F(9, 109.996)=20.644$, $p<.0001$<br>Outcome $F(1, 20.878)=24.739$, $p<.0001$<br>Interaction $F(6, 110.337)=5.431$, $p<.0001$ | Day 2: ABA-S > ABA-R $p=.014$<br>Days 3-6: ABA-S > ABA-R all $ps<.0001$ |
| **3C** | Two-way RM ANOVA<br><u>ONLY INCLUDES ABA-S and ABA-R</u> | Time $F(23, 391)=18.93$, $p<.0001$<br>Outcome $F(1, 17)=13.06$, $p=.0021$<br>Interaction $F(23, 391)=3.574$, $p<.0001$ | 1800h: ABA-S > ABA-R $p=.0003$<br>1900h: ABA-S > ABA-R $p<.0001$<br>2000h: ABA-S > ABA-R $p=.0036$<br>2100h: ABA-S > ABA-R $p=.0063$<br>0900h: ABA-S > ABA-R $p=.0156$ |
| **3E** | Two-way RM ANOVA | Time $F(23, 391)=22.33$, $p<.0001$<br>Outcome $F(1, 17)=0.1744$, $p=.6814$<br>Interaction $F(23, 391)=3.433$, $p<.0001$ | 1300h: ABA-R > ABA-S $p<.0001$ |
| **3F** | Two-way RM ANOVA | Phase $F(1, 24)=204.3$, $p<.0001$<br>Group $F(2, 24)=11.66$, $p=.0003$<br>Interaction $F(2, 24)=11.33$, $p=.0003$ | Phase: all groups ABA > Baseline $p<.0001$<br>Group: ABA-S > ABA-R $p=.0004$; ABA-S > RW $p=.0038$<br>ABA phase: ABA-S > ABA-R $p<.0001$; ABA-S > RW $p<.0001$ |
| **3G** | One-way ANOVA | $F(2, 24)= 11.33$, $p=.0003$ | ABA-S > ABA-R $p=.0022$<br>ABA-S > RW $p=.0008$ |
| **3H** | Linear regression | ABA-S: $r=.4970$, $R^2=.2470$, $p=.0002$<br>ABA-R: $r=.1539$, $R^2=.02367$, $p=.1476$<br>RW: $r=.3553$, $R^2=.1263$, $p=.0012$ | |
| **3I** | Linear regression | ABA-S: $r=.5574$, $R^2=.3107$, $p<.0001$<br>ABA-R: $r=.5508$, $R^2=.3034$, $p<.0001$<br>RW: $r=.1986$, $R^2=.03945$, $p=.0774$ | |
| **3J** | Two-way RM ANOVA | Phase $F(1, 24)=146.6$, $p<.0001$<br>Group $F(2, 24)=5.870$, $p=.0084$<br>Interaction $F(2, 24)=14.84$, $p<.0001$ | Phase: ABA-S $p=.0007$; ABA-R $p<.0001$; RW $p<.0001$<br>Group: ABA-R > ABA-S $p=.0093$; ABA-R > RW $p=.0430$<br>ABA phase: ABA-R > ABA-S $p<.0001$; ABA-R > RW $p=.0002$ |
| **3K** | One-way ANOVA | $F(2, 24)= 14.84$, $p<.0001$ | ABA-R > ABA-S $p<.0001$<br>ABA-R > RW $p=.0019$ |
| **3L** | Linear regression | ABA-S: $r=.2823$, $R^2=.07969$, $p=.0666$<br>ABA-R: $r=.4569$, $R^2=.2088$, $p<.0001$<br>RW: $r=.0957$, $R^2=.009167$, $p=.4237$ | |
| **3M** | Linear regression | ABA-S: $r=.0139$, $R^2=.0007975$, $p=.9293$<br>ABA-R: $r=.4876$, $R^2=.2378$, $p<.0001$<br>RW: $r=.3484$, $R^2=.1214$, $p=.0027$ | |

**Supplementary Table 3.** Statistical test details and results for data presented in Figure 4.

| **Figure** | **Statistical test** | **Main analysis result** | **Post-hoc multiple comparisons** |
| --- | --- | --- | --- |
| **4B** | Two-way RM ANOVA | Zone $F(3, 68)=111.2$, $p<.0001$<br>ABA outcome $F(1, 68)=7.200e-006$, $p=.9979$<br>Interaction $F(3, 68)=0.08631$, $p=.9673$ | |
| **4C** | Two-way RM ANOVA | Zone $F(3, 68)=155.5$, $p<.0001$<br>ABA outcome $F(1, 68)=8.307$, $p=.0053$<br>Interaction $F(3, 68)=0.9624$, $p=.4156$ | Centre zone: ABA-S > ABA-R $p=.0355$ |
| **4D** | Independent samples t-test | $t(17)=1.630$, $p=.1215$ | |
| **4E** | Independent samples t-test | $t(17)=0.5274$, $p=.6048$ | |
| **4F** | Independent samples t-test | $t(17)=1.138$, $p=.2709$ | |
| **4G** | Linear regression | ABA-S: $r=.6495$, $R^2=.4219$, $p=.0421$<br>ABA-R: $r=.2141$, $R^2=.04586$, $p=.5801$<br>Combined ABA: $r=.1946$, $R^2=.03785$, $p=.4248$ | |
| **4I** | Independent samples t-test | $t(17)=0.1753$, $p=.8629$ | |
| **4J** | Independent samples t-test | $t(17)=0.5757$, $p=.5724$ | |
| **4K** | Independent samples t-test | $t(17)=0.4199$, $p=.6799$ | |
| **4L** | Independent samples t-test | $t(17)=0.2738$, $p=.7875$ | |
| **4M** | Linear regression | ABA-S: $r=.7706$, $R^2=.5938$, $p=.0091$<br>ABA-R: $r=.2660$, $R^2=.07075$, $p=.4891$<br>Combined ABA: $r=.2637$, $R^2=.06955$, $p=.2753$ | |
| **4N** | Linear regression | ABA-S: $r=.4866$, $R^2=.2368$, $p=.1538$<br>ABA-R: $r=.2435$, $R^2=.05931$, $p=.5278$<br>Combined ABA: $r=.1152$, $R^2=.01326$, $p=.6387$ | |
| **4P** | Independent samples t-test | $t(17)=2.060$, $p=.0551$ | |
| **4Q** | Linear regression | <u>Baseline</u><br>ABA-S: $r=.1793$, $R^2=.03216$, $p=.6201$<br>ABA-R: $r=.4835$, $R^2=.2338$, $p=.1873$<br>Combined ABA: $r=.1954$, $R^2=.03819$, $p=.4227$<br><u>ABA</u><br>ABA-S: $r=.3607$, $R^2=.1301$, $p=.3059$<br>ABA-R: $r=.4800$, $R^2=.2304$, $p=.1909$<br>Combined ABA: $r=.0430$, $R^2=.001853$, $p=.8611$ | |
| **4R** | Linear regression | ABA-S: $r=.5102$, $R^2=.2603$, $p=.1319$<br>ABA-R: $r=.7147$, $R^2=.5108$, $p=.0305$<br>Combined ABA: $r=.5205$, $R^2=.2709$, $p=.0223$ | |

**Supplementary Table 4.** Statistical test details and results for data presented in Figure 5.

| Figure | Statistical test | Main analysis result | Group size |
| --- | --- | --- | --- |
| **5B1** | Welch's One-way RM ANOVA | $W(4, 12.27)=1.299, p=.3238$ | RANTES: ABA-S n=6, ABA-R n=6, FR n=6, RW n=7, CON n=7 |
| **5B2** | Welch's One-way RM ANOVA | $W(4, 12.34)=0.8452, p=.5223$ | |
| **5B3** | Welch's One-way RM ANOVA | $W(4, 12.54)=1.077, p=.4086$ | |
| **5C1** | Welch's One-way RM ANOVA | $W(4, 11)=2.172, p=.1394$ | IL-4: ABA-S n=6, ABA-R n=6, FR n=3, RW n=6, CON n=6 |
| **5C2** | Welch's One-way RM ANOVA | $W(4, 8.492)=0.7422, p=.5880$ | |
| **5C3** | Welch's One-way RM ANOVA | $W(4, 10.69)=3.202, p=.0581$ | |
| **5D1** | Welch's One-way RM ANOVA | $W(4, 10.57)=1.218, p=.3599$ | IL-6: ABA-S n=6, ABA-R n=6, FR n=6, RW n=6, CON n=5 |
| **5D2** | Welch's One-way RM ANOVA | $W(4, 10.63)=0.3845, p=.8152$ | |
| **5D3** | Welch's One-way RM ANOVA | $W(4, 10.52)=1.132, p=.3929$ | |
| **E** | Linear regression | ABA-S: $r=.4362, R^2=.1903, p=.4627$<br>ABA-R: $r=.9128, R^2=.8332, p=.0305$<br>Combined ABA: $r=.5017, R^2=.2517, p=.1396$ | IL-6: ABA-S n=5, ABA-R n=5, Combined ABA n=10 |
| **F** | Linear regression | ABA-S: $r=.8947, R^2=.8004, p=.0404$<br>ABA-R: $r=.8978, R^2=.8061, p=.0386$<br>Combined ABA: $r=.3927, R^2=.1542, p=.2617$ | |
| **G** | Linear regression | ABA-S: $r=.9513, R^2=.9049, p=.0128$<br>ABA-R: $r=.0614, R^2=.003770, p=.9219$<br>Combined ABA: $r=.3021, R^2=.09124, p=.3963$ | |
| **H** | Linear regression | ABA-S: $r=.8907, R^2=.7934, p=.0173$<br>ABA-R: $r=.1198, R^2=.01436, p=.8211$<br>Combined ABA: $r=.1201, R^2=.01442, p=.7101$ | IL-4: ABA-S n=6, ABA-R n=6, Combined ABA n=12 |

**Supplementary Figures 1-3.**
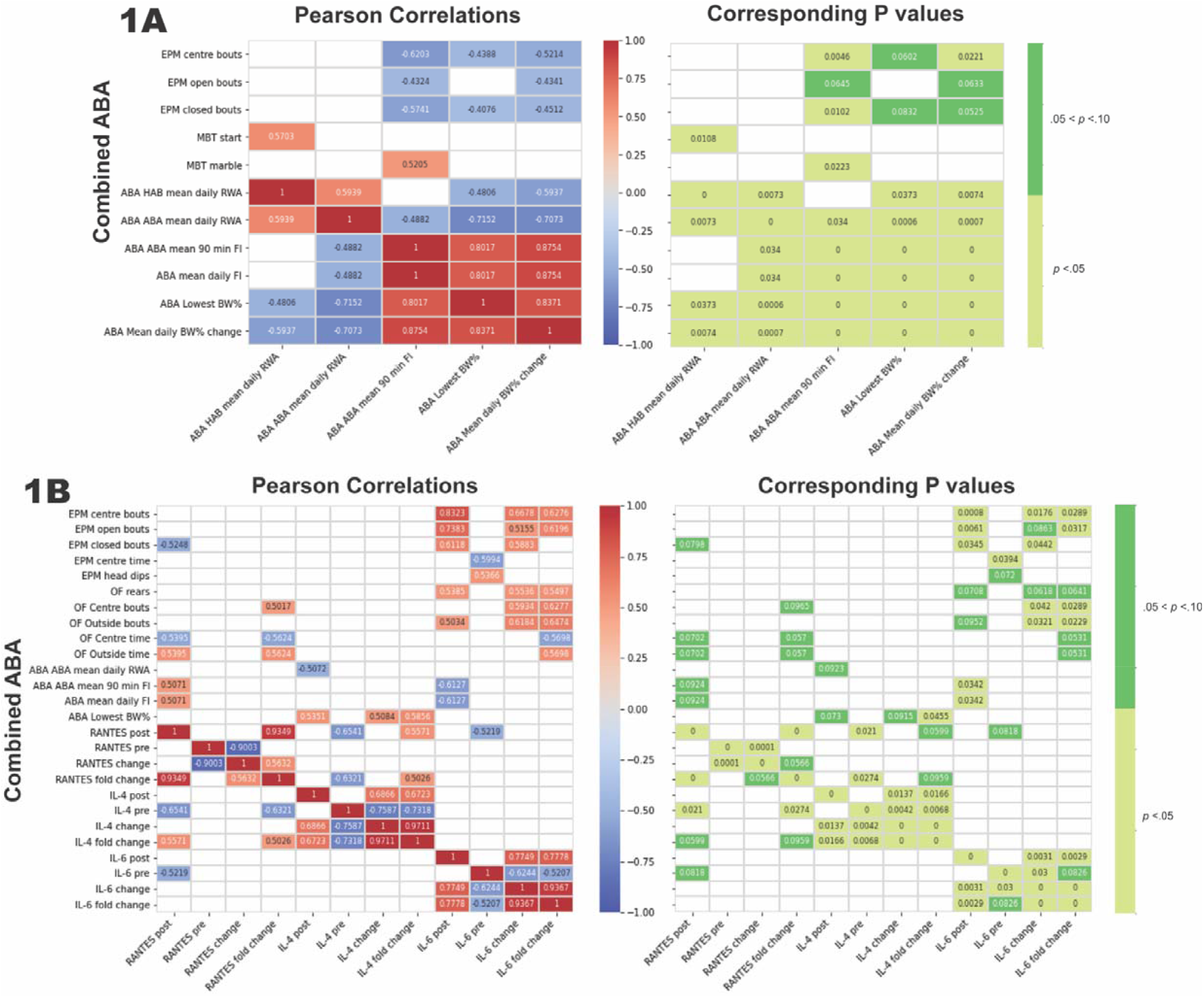
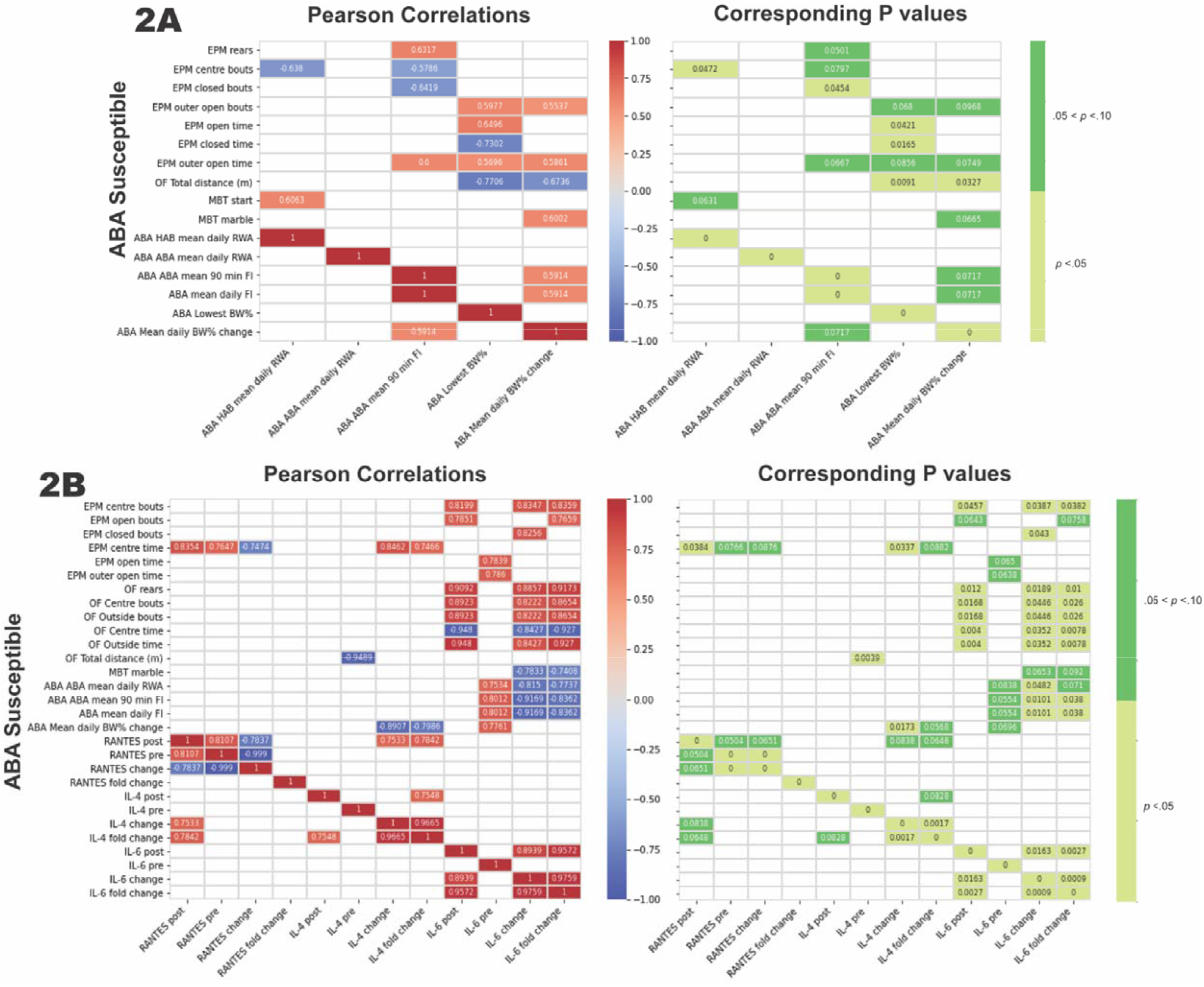
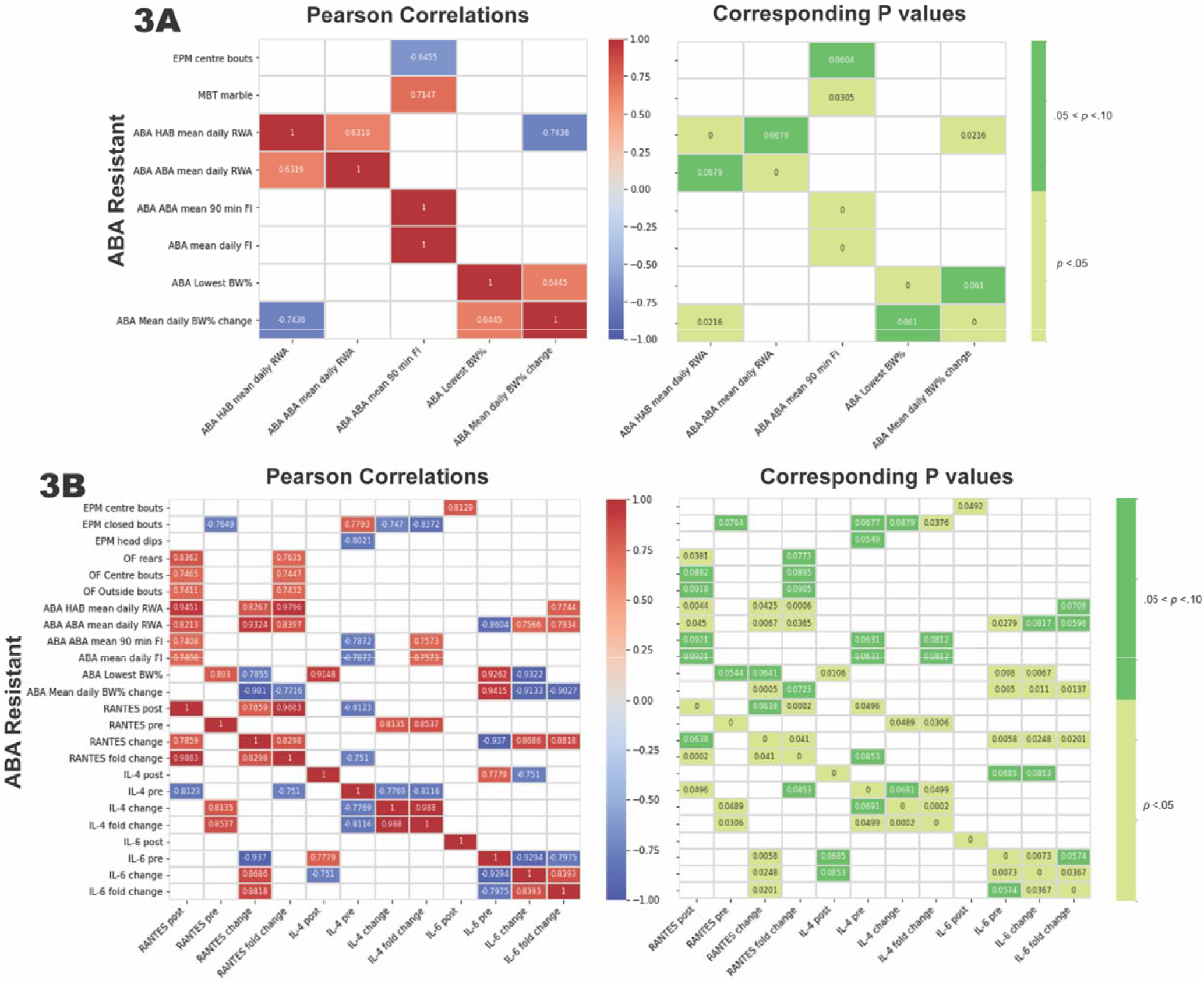
For each of the **1)** Combined ABA, **2)** ABA Susceptible, and **3)** ABA Resistant groups, Python was used to generate Pearson’s correlation and corresponding *p* value matrices for **A)** outcomes from behavioural tests prior to ABA vs. ABA outcome measures, and **B)** outcomes from behavioural tests prior to ABA and ABA outcome measures vs. cytokine levels (pre-exposure, post-exposure, absolute change and fold-change from pre- to post-exposure). Python was then utilized to produce the above plots of these matrices showing data only for pairs of variables where *p*<.01. Group sizes for behavioural correlations: Combined ABA *n=*19; ABA-S *n=*10; ABA-R *n=*9 and for cytokine correlations: Combined ABA *n=*12; ABA-S *n=*6; ABA-R *n=*6. Abbreviations: EPM, elevated plus maze; OF, open field; bouts, number of entries into zone; time, percentage of total 10-minute test duration, MBT, marble burying test; start, latency in seconds until first episode of digging; marbles, number of interactions with marbles; ABA, activity-based anorexia; ABA HAB, baseline phase of ABA; ABA ABA, ABA (food restriction) period of ABA; RWA, running wheel activity; FI, food intake; BW%, body weight %.

